# Synapses, predictions, and prediction errors: a neocortical computational study of MDD using the temporal memory algorithm of HTM

**DOI:** 10.1101/2022.06.29.498015

**Authors:** Mohamed A. Sherif, Mostafa Z. Khalil, Rammohan Shukla, Joshua C. Brown, Linda L. Carpenter

## Abstract

**Background:** Synapses and spines are central in major depressive disorder (MDD) pathophysiology, recently highlighted by ketamine’s and psilocybin’s rapid antidepressant effects. The Bayesian brain and interoception perspectives formalize MDD as being “stuck” in affective states constantly predicting negative energy balance. We examined how synaptic atrophy relates to the predictive function of the neocortex and thus to symptoms, using temporal memory (TM), an unsupervised machine-learning algorithm. TM represents a single neocortical layer, learns in real-time using local Hebbian-learning rules, and extracts and predicts temporal sequences.

**Methods:** We trained a TM model on random sequences of upper-case alphabetical letters, representing sequences of affective states. To model depression, we progressively destroyed synapses in the TM model and examined how that affected the predictive capacity of the network.

**Results:** Destroying 50% of the synapses slightly reduced the number of predictions, followed by a marked drop with further destruction. However, reducing the synapses by 25% dropped the confidence in the predictions distinctly. So even though the network was making accurate predictions, the network was no longer confident about these predictions.

**Conclusions:** These findings explain how interoceptive cortices could be stuck in limited affective states with high prediction error. Growth of new synapses, e.g., with ketamine and psilocybin, would allow representing more futuristic predictions with higher confidence. To our knowledge, this is the first study to use the TM model to connect changes happening at synaptic levels to the Bayesian formulation of psychiatric symptomatology, making it possible to understand treatment mechanisms and possibly, develop new treatments.

**Graphical abstract:** 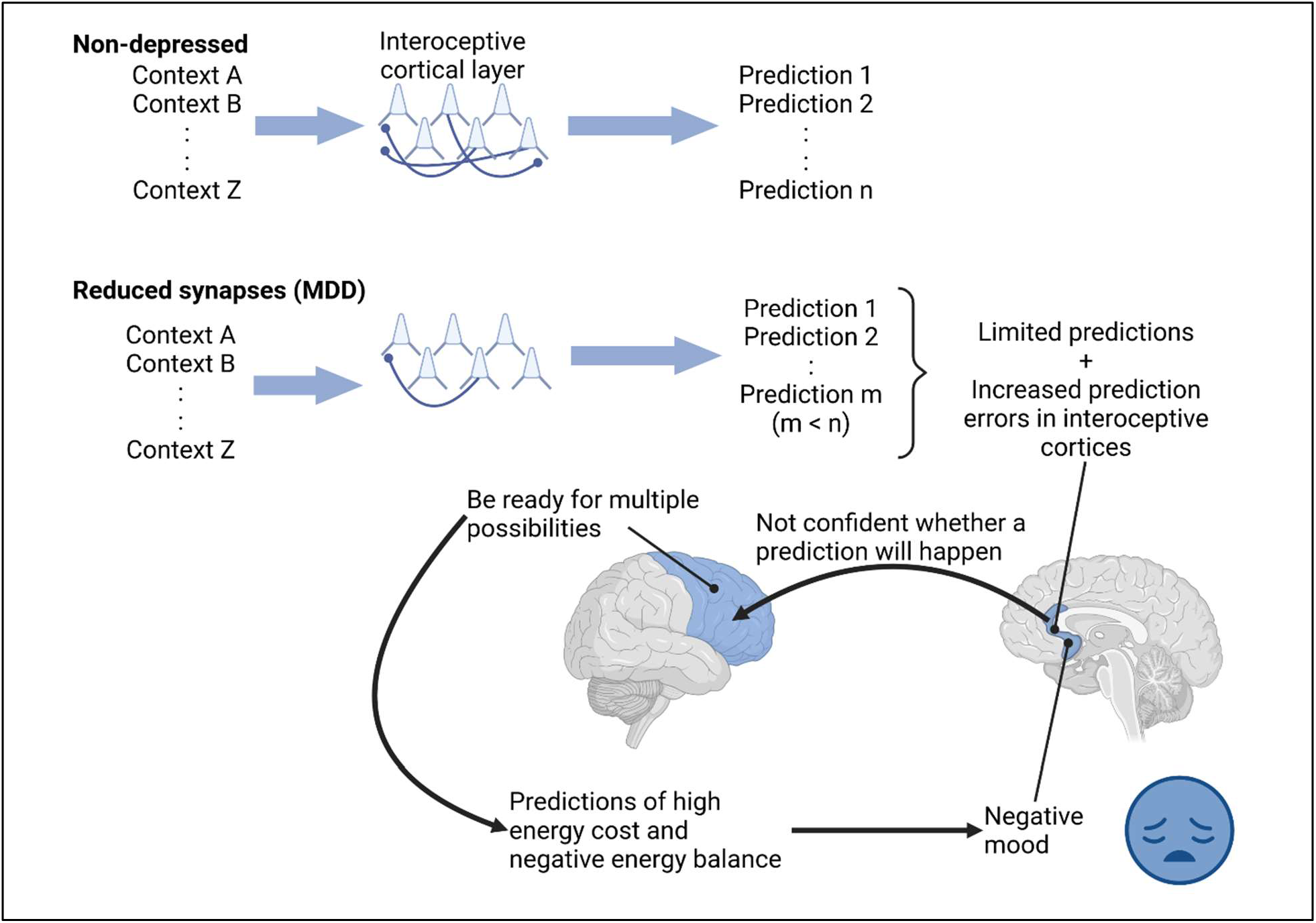

## Introduction

Synapses and spines play an integral role in the pathophysiology of major depressive disorder (MDD), a severely disabling disease that affects many aspects of life (1–3). Postmortem evidence shows a reduction in spine density (4) and decreased synapse numbers in the dorsolateral prefrontal cortex (DLPFC) in patients diagnosed with MDD (5). Paired associative stimulation using transcranial magnetic stimulation (TMS) showed impaired plasticity in patients struggling with MDD, suggesting abnormal synaptic functioning (6). In chronic stress animal models of MDD, dendritic spines of layer 2/3 in the medial prefrontal cortex are lost (7–9). Chronic imipramine or fluoxetine restored spine loss in both medial prefrontal cortex layer 2/3 pyramidal neurons and hippocampal CA3 in animal models of chronic stress (10). Likewise, a single session of repetitive TMS in hippocampal CA1 slice cultures increased spine size and functional insertion of the GluA1 subtype of AMPA receptors consistent with long-term potentiation (LTP) of excitatory synapses (11). Cambiaghi et al (12) showed increased total spine density in apical and basal dendrites of pyramidal neurons layer 2/3, with a predominance of thin spines following 5 days of high frequency repetitive TMS applied to motor cortex. The treatment with TMS also increased dendritic complexity. These findings may be extrapolated to healthy humans as suggested by LTP-like changes after a single-session of repetitive TMS in the motor cortex when combined with pharmacologic n-methyl-d-aspartate (NMDA) receptor agonism (13, 14). Similar effects on spine maturation have been reported with electroconvulsive therapy in animal models (15, 16).

Single doses of ketamine (17–21) and psilocybin (22) result in rapid antidepressant effects in depressed patients that appear to be mediated by restoring synapses and spines. A single dose of ketamine increased the rate of spine formation in non-stressed animals (23). Following chronic stress, ketamine restored the number of neocortical synapses in animal models in layer 5 of the medial prefrontal cortex (24, 25). A single dose of ketamine also protected spine loss induced by chronic stress (26). The deleterious effects on spines were mediated by proteins like REDD1 that belong to the mTOR pathway in layer 5 apical dendrites, which increased with chronic stress. REDD1 was also elevated in the prefrontal cortex in patients diagnosed with MDD (27). An increase in spine densities suggests the formation of more synapses (28), and mediated synchronized firing of ensembles encoding behavioral manifestations affected by depression (29). Local infusion of ketamine into the infralimbic cortex of rats resulted in the reversal of depression-like behavior. This effect was also generated by optogenetic stimulation of layer 5 pyramidal neurons and resulted in similar increases in spine density in apical tufts (30). Related findings by Shao et al. showed that psilocybin increased dendritic spines on both apical and basal dendrites in the medial frontal cortex in a mouse model of depression, reversing the behavioral manifestations of depression (31). Together these clinical and preclinical findings converge to illustrate a critical role for synaptic function and number in the pathology of depression and in antidepressant treatment effects, but the direct and dynamic investigation of neuronal synapses in living clinical samples is not currently possible.

To bridge synaptic changes to symptomatology in MDD, it is helpful to consider the function of the central nervous system from an evolutionary perspective. The brain has evolved to optimize energy consumption as the body acts and grows, a phenomenon termed allostasis, which means stabilizing in the face of change (32–35). Instead of reacting to inputs, the neocortex develops an internal model of the world and of the body, and uses the model to predict input changes and update the model predictions based on mismatches with the inputs. This has been formulated into the Bayesian brain and predictive coding frameworks (36, 37).

In the motor or “action” realm, instead of minimizing prediction error by updating the model to match inputs, the body minimizes the prediction error by enacting the prediction. For example, before a person reaches out to grasp a cup of water, the motor cortex generates predictions about the action, such as “what would I experiences when I am grasping that cup of water I see in front of me on the counter?” Relevant visual, tactile, and proprioceptive predictions are projected to the spinal cord. Spinal cord reflex arcs then “enact” or fulfill these predictions by moving the person’s hand and grasping the cup of water, thus, reducing prediction error. This is termed active inference (38).

Emotional states and their associated autonomic, hormonal, and immune system changes are hypothesized to reflect enactments of the brain’s predictions based on inputs from the inner milieu of the body, as well as from the external world (interoceptive predictions). Formulated in the Embodied Predictive Interoception Coding (EPIC) framework, mood has been considered a low-dimensional summary of the predictions related to these inputs (39). Interoceptive predictions occur in interoceptive limbic cortical areas, such as the anterior cingulate cortex (ACC) and insula, the areas affected in chronic stress models of depression where ketamine was found to be helpful in animal models. Applying the interoceptive theory to understand states of clinical depression, Barrett et al. hypothesized that the brain is locked-in, i.e., stuck in a “metabolically-inefficient internal model of the body in the world” (32, 40). Ruminations could manifest such locked-in states, where the brain keeps coming back to the same thought content. Such states could arise as a consequence of the abnormalities in interoceptive cortical areas that have been revealed by neuroimaging studies in depressed samples (41), and suggested as sites for DBS stimulation treatment for MDD (42). Indeed, TMS has better outcomes when post-hoc modeling shows DLPFC stimulation has strong functional connectivity (negative correlation) with the ACC (43, 44). Taken further, personalized targeting to the ACC was one of several changes that produced the strongest TMS outcomes to date (45), and may become the standard for TMS therapy (46).

Machine learning algorithms constrained by neocortical neurobiology can connect synaptic atrophy to impaired predictions, a functional construct more proximal to symptoms. The neocortex is hypothesized to be a spatio-temporal pattern-predictive machine (47–49). The ability of the neocortex to learn from new information on the fly and adjust its predictions accordingly is an integral part of this process (50). This is referred to as “online learning” in the machine learning literature and is different from batch training, where an algorithm learns a training set, then is used on a different set. We used the temporal memory (TM) algorithm, a computer model of neocortical layers, to investigate the link between synaptic changes and prediction making. TM is one algorithm of the hierarchical temporal memory (HTM) framework. The underlying assumption of HTM is that different neocortical regions implement versions of the same cortical algorithm (51) for online unsupervised sequence learning and prediction (52, 53). Multiple features of the neocortex constrain HTM algorithms in ways unlike most of the current artificial neural networks (54). For example, the units or “neurons” in the HTM algorithms exist in one of three states: active, inactive, and predictive. Neurobiologically, the predictive state of a neuron maps to subthreshold depolarization of the soma by dendritic NMDA spikes, making it easier for the neuron to fire. NMDA spikes are triggered by the co-activation of synapses clustered together on a dendritic segment, indicating the detection of a specific input pattern (55). HTM exhibits online learning, where the algorithm continuously learns and predicts, rather than being trained in batches like artificial neural networks. In addition, the HTM algorithms learn by local unsupervised Hebbian-learning rules similar to synaptic plasticity that does not require error back propagation. HTM algorithms act on sparse-distributed representations (SDR) of data, which takes the stochastic nature of synaptic activation into account (52).

With a TM model representing a neocortical layer of neurons that learns and predicts sequences of inputs, this study explored a possible connection between having a reduced number of synapses (i.e., characteristic of MDD) and being locked in dysfunctional predictive states experienced as negative affective states. The TM model we use contains basal dendrites, which provide temporal context to the presented stimuli (52, 54). We ran simulations to examine how many predictions are generated given a particular context, and then examined how synaptic loss affected the accuracy and confidence of the predictions made.

## Methods

### Inputs, temporal memory (TM) models, and training

The input patterns consisted of an array of ten randomly generated sequences of 11 upper-case letters (A to Z) (Fig. 1). A TM model is composed of multiple minicolumns. Each upper-case letter was represented in the TM network by activating neurons in 20 consecutive minicolumns, resulting in a network with 520 minicolumns (20 minicolumns x 26 upper-case letters). Each minicolumn contained 32 neurons.

**Fig. 1:**
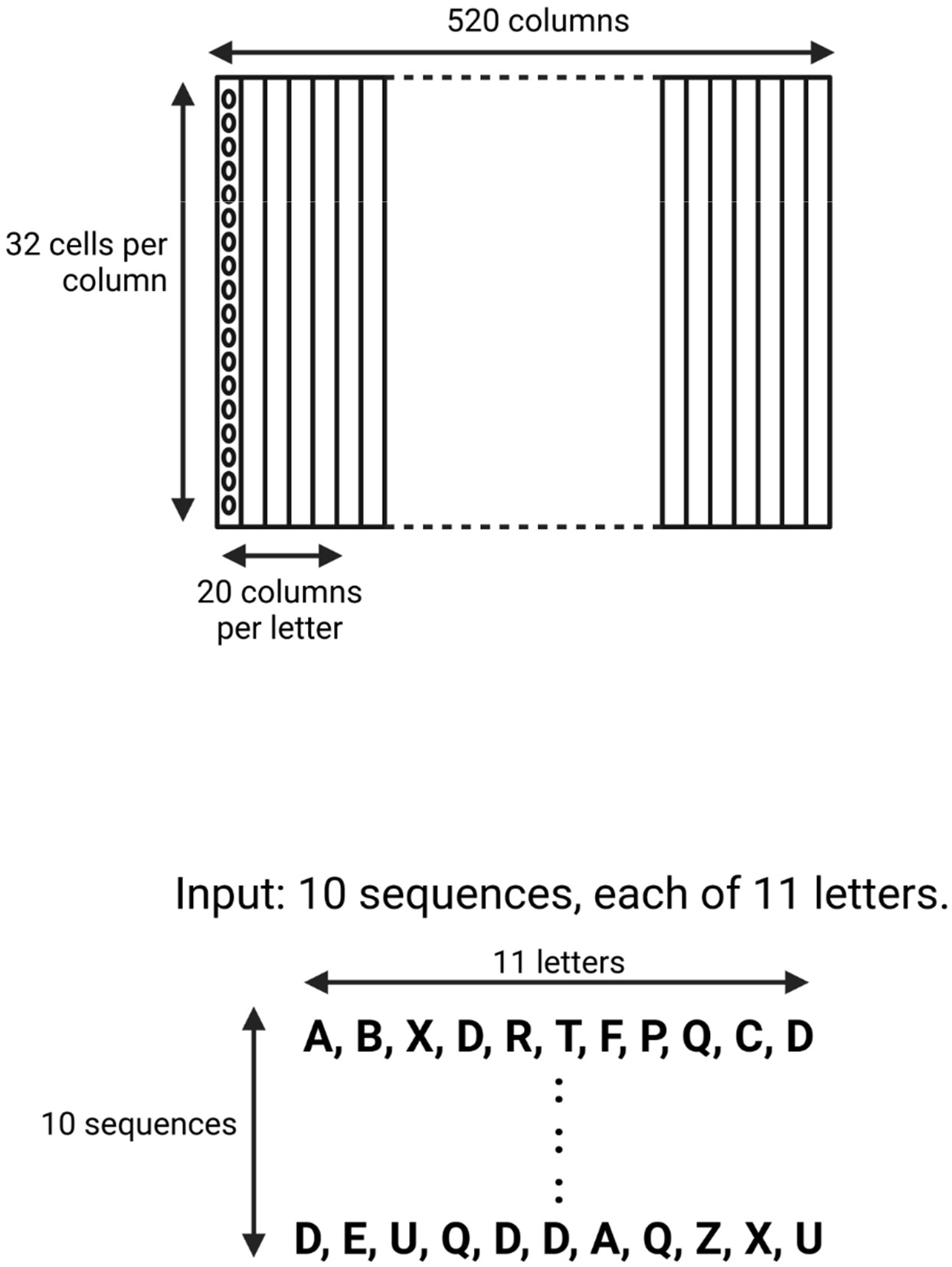
Schematic of the neocortical layer temporal memory (TM) model and the input it learned. The network consisted of 520 columns, with 32 cells per column. Each upper-case letter has a receptive field of 20 columns. The input consists of 10 sequences, each of 11 upper-case letters selected randomly from a uniform distribution with replacement.

Using the Numenta Platform for Intelligent Computing (NuPIC) package (version 1.0.5) we generated five temporal memory (TM) models (parameters in Table 1) using five different randomization seeds. We then trained each model on five different versions of the input patterns. The training steps are explained in detail in the results section and in schematic Fig. 2. On average, each TM model correctly learned each sequence of the 11 letters over ten repeated presentations. Therefore, for each of the five versions of the input pattern, we ran 1100 iterations of simulations to ensure the model had learned the full input pattern. This generated five trained network models for each input pattern, resulting in 25 trained TM models. Next, we calculated the total number of future predictions a network can make by summing the number of predictions for each of the upper-case letters:

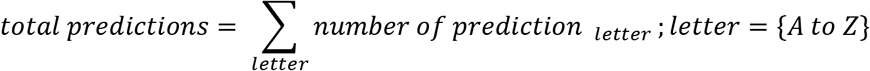

**Fig. 2:**
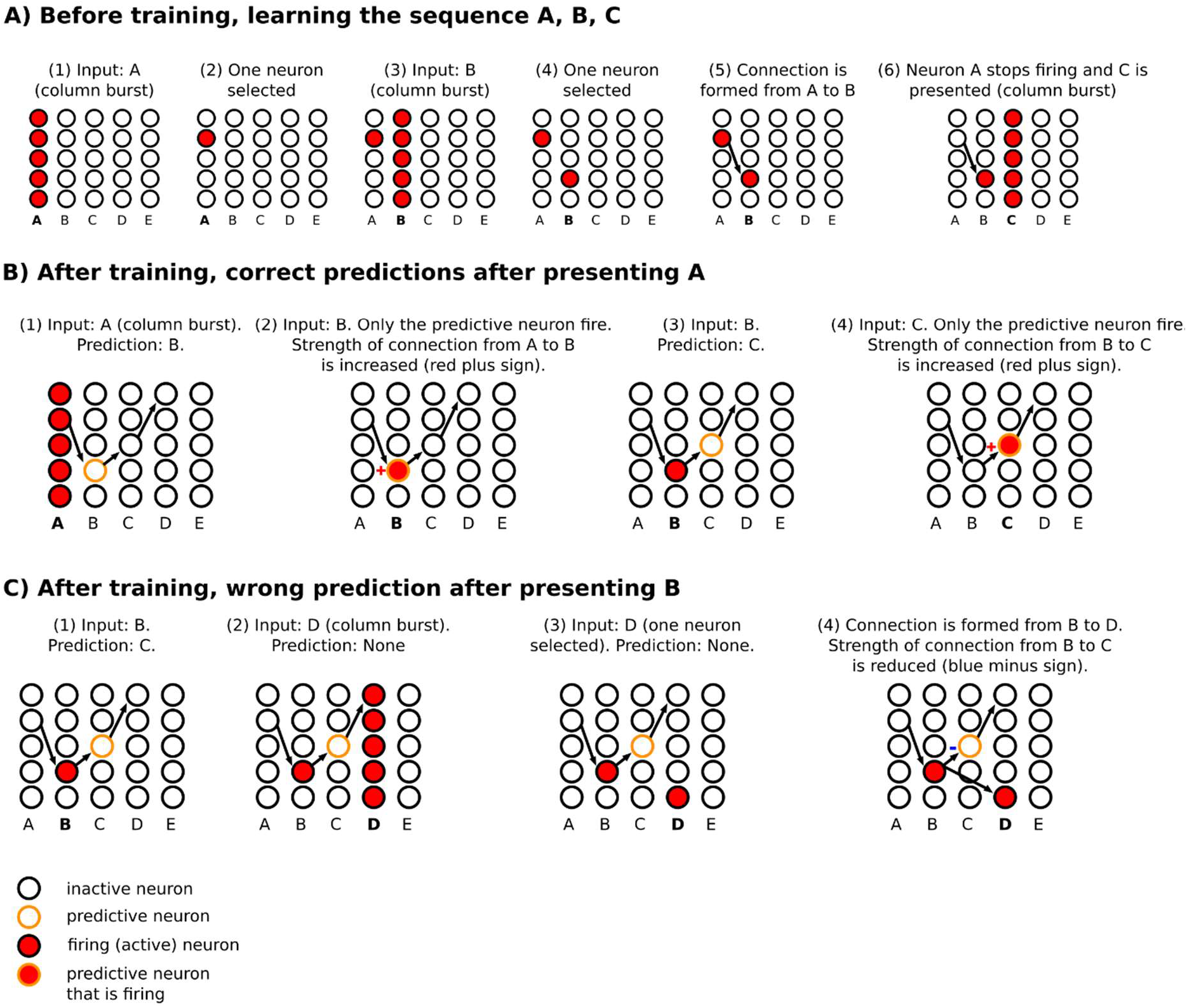
Schematic diagram of steps for learning and predicting in HTM. **(A)** The sequence A, B, C is presented to the temporal memory algorithm. Each minicolumn represents the receptive field of an input. Since the network has not learned any sequences, the presentation of each of the letters fires all the neurons in a minicolumn (minicolumn burst). **(B)** After learning, the presentation of A predicts B. Presenting B confirms the prediction, strengthening the connection from A to B (same for predicting C after presenting B). **(C)** After B is presented, C is predicted. However, D is presented after that, firing all the neurons in minicolumn corresponding to D. This results in making a new connection from a neuron in the B minicolumn to a neuron in the D minicolumn, with reduction of connection strength from B to C. Of note, we are separating the before training and after training phase to simplify the description, but learning and predicting happens continuously.

### Destroying synapses

Using five randomization seeds that were different from the ones used to generate the TM models, we progressively removed fractions of synapses, i.e., we destroyed 25%, then 50%, then 60%, then 70%, and finally 80% of the total number of synapses. For each of the 25 networks (5 networks x 5 random variations of input sequences), this step created five different versions of “depressed” networks for each percentage of destroyed synapses, resulting in 125 versions of networks for each percentage of destroyed synapses. We then calculated the total number of predictions made at each level of synaptic destruction.

### Calculating prediction error

As described in more detail in the results, when a minicolumn receives a predicted input, only the neurons in the predictive state fire and they are the winner cells. But when a minicolumn receives input, and none of the cells in that minicolumn were in a predictive state, all the neurons in the minicolumn fire (minicolumn burst); in this case, winner cells are selected at random from the active neurons. To estimate the prediction error, we calculated the ratio of winner cells to active cells:

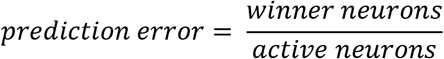

Lower prediction error represents a better match between prediction and subsequent input. Higher prediction error, usually due to a higher number of active cells when a minicolumn bursts, means a higher mismatch occurred between the prediction and subsequent input.

## Results

### The network learned and predicted accurately with training

We presented five input arrays to each of five temporal memory (TM) versions. Each input array consisted of 10 rows x 11 minicolumns of randomly-selected upper case letters (A to Z) from a uniform distribution. It took each TM network an average of 1100 iterations to learn to predict the input patterns with 100% prediction accuracy (Fig. 3A). For clarity, below we describe separately how the learning took place before and after training. However, the TM learns and predicts continuously.

**Fig. 3:**
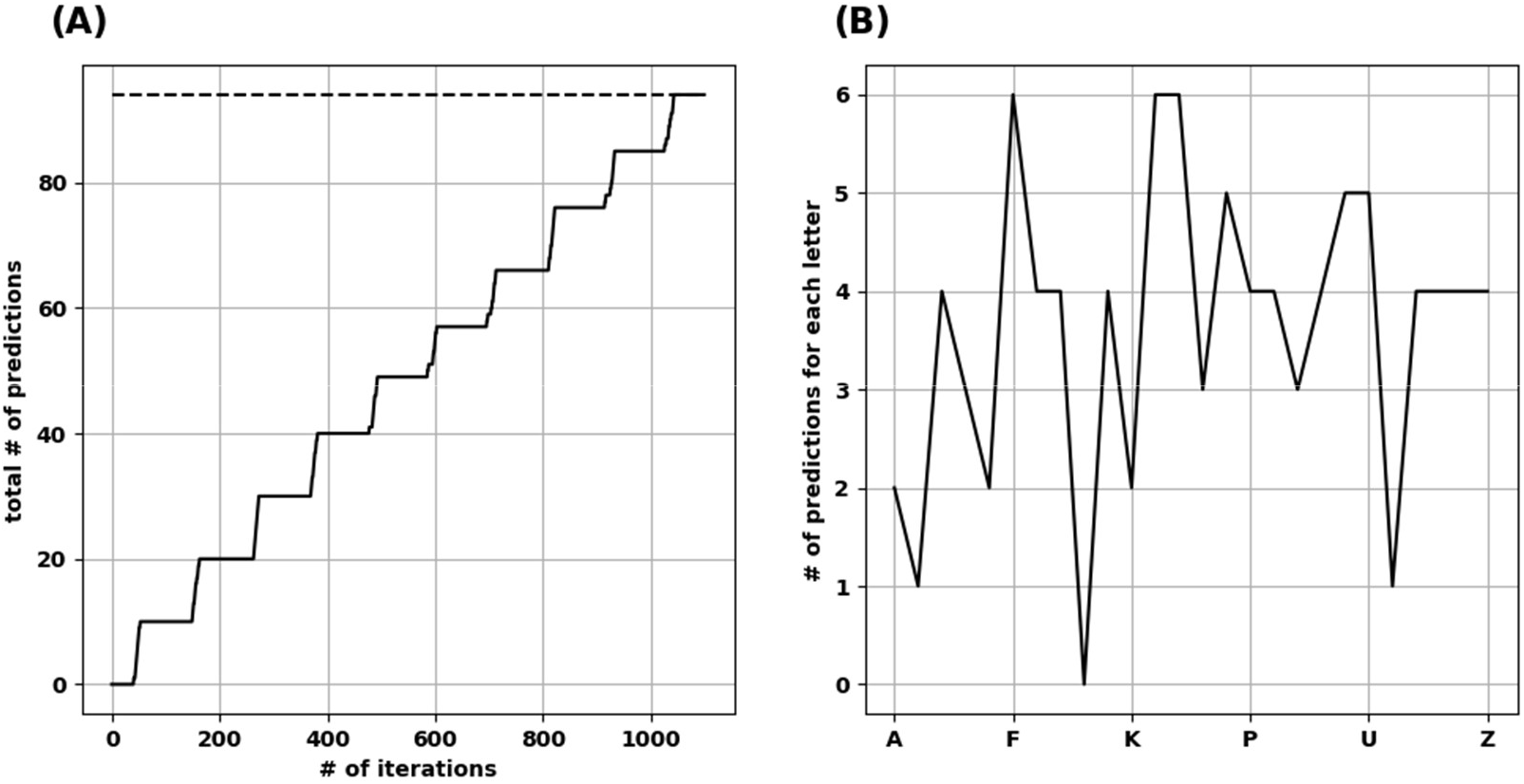
The network learned and reached the number of ideal predictions. **(A)** Over 1100 iterations, the network progressively made predictions approaching ideal predictions (dotted line). **(B)** Number of predictions when the network was presented by each uppercase letter, resulting in a “prediction landscape”.

#### Before training

The basic learning steps have been described in multiple papers (52, 56, 57) and are illustrated schematically in Fig. 2A. In summary, the network learned as follows:

1. Before any learning, no predictions were made as the neocortical layer was learning the sequences for the first time.
2. When a stimulus was presented (e.g., “A”), all neurons in the minicolumns corresponding to the presented letter, i.e., its receptive fields, were activated.
3. Winner neurons in the first set of minicolumns were randomly selected (one neuron from each column).
4. With the presentation of the following sequence (e.g., “B”), all neurons in the receptive field minicolumns were activated as well.
5. Winner neurons in the second set of minicolumns were randomly selected.
6. A connection was made from the winner neurons of the previous time step to the winner neurons in the current time step.
7. Repeat from step number 2 until the sequence ended.

#### After training

After a network learned the sequences of letters, the network continued predicting and updating its predictions as follows (Fig. 2B):

1. After the first stimulus was presented, since it was the first one, there were no predictions before it. All neurons in the receptive field minicolumns fired.
2. The firing of the neurons across the receptive field set their target neurons into a predictive state. Biologically, this occurs through activation of an NMDA spike that depolarizes the soma membrane to sub-threshold voltage (55, 58). Through learning, the neurons in the predictive state were in minicolumns corresponding to the next predicted letter.
3. When the subsequent input arrived, the next step varied depending on whether the input matched the predicted input, as follows.

a. If the input matched the minicolumns that contained the predictive neurons, then within each minicolumn, the predictive neurons fired earlier than the other neurons in the minicolumn because they were closer to the spiking threshold. This firing in turn inhibited the other neurons in the minicolumn (winner-takes-all), and the predictive neurons were selected as the winner neurons. The synapses between the neurons that were active in the previous time step and the predictive-turned-winner neurons will be strengthened in a Hebbian manner, where neurons that fire repeatedly in close temporal proximity to each other have stronger connections (59).
b. If the input did *not* match the minicolumns that contained the predictive neurons, all neurons in each minicolumn corresponding to the current input were activated and the winner neurons were randomly selected. New synapses were then formed between the neurons that were active in the previous time step and the randomly-selected winner neurons in the current time step. Also, the synapses that were projecting to the incorrectly predicted neurons were weakened. The prediction error signal in these cases was the activation of all neurons in the minicolumns receiving the input.

In the TM model, formation of synaptic connections and changes in their strength is not done by changing the synaptic weights (magnitude of synaptic influence on the neuron). Instead, it is done by modifying synaptic permanence. Permanence is how resistant the synapses are for removal, where stronger synapses are more resistant to atrophy (52).

### The total number of predictions a TM can make

To calculate the total number of predictions each trained TM version is capable of making (with all synapses intact), we presented each uppercase alphabetical letter to the network while pausing new learning to avoid changes in synaptic plasticity. We calculated how many letters were predicted on the next time step, to create a “prediction landscape” (Fig. 3B). We compared each prediction landscape to the number of letter dyads, and as expected, we found no difference between a well-trained network and the frequency of letter dyads in the presented sequences. Summing the predictions across the upper-case letters resulted in the total number of predictions each trained TM makes. The mean number of total predictions across the 25 TM versions was 94.6 (std. dev. = 1.85).

### Loss of predictions and increased prediction error with synaptic destruction

To understand how synaptic atrophy can affect the predictive function of a neocortical layer, we randomly destroyed the number of synapses in fractions (25%, 50%, 60%, 70%, and 80%) across all trained TM versions (5 TM versions trained across 5 variations of input sequences) and examined the number of predictions made by the models. Reducing the number of synapses resulted in a progressively flatter prediction landscape, starting at the point of 50% synapse destruction and reaching a flat line of zero predictions when there were 20% remaining synapses (Fig. 4A). To ensure the reproducibility of this result, we used five random seeds to randomly select the synapses to be destroyed for each fraction of synaptic removal, resulting in 125 network versions for each fraction removed. With 75% of synapses intact, the mean number of predictions was the same as it was at baseline when all synapses were present (Fig. 4B, black data points and left y-axis). At 50% removal, the mean number of predictions decreased slightly to 89.1 (std. dev. = 2.7), which is still close to the number of predictions a “healthy” network (100% synapses) had. With 40% of synapses intact, the mean number of predictions dropped to 34.9 (std. dev. = 2.97). With 30% of synapses remaining, the average number of predictions was 2.5 (std. dev. = 1.7), and when only 20% of synapses were remaining, no predictions were generated across the 125 networks.

**Fig. 4:**
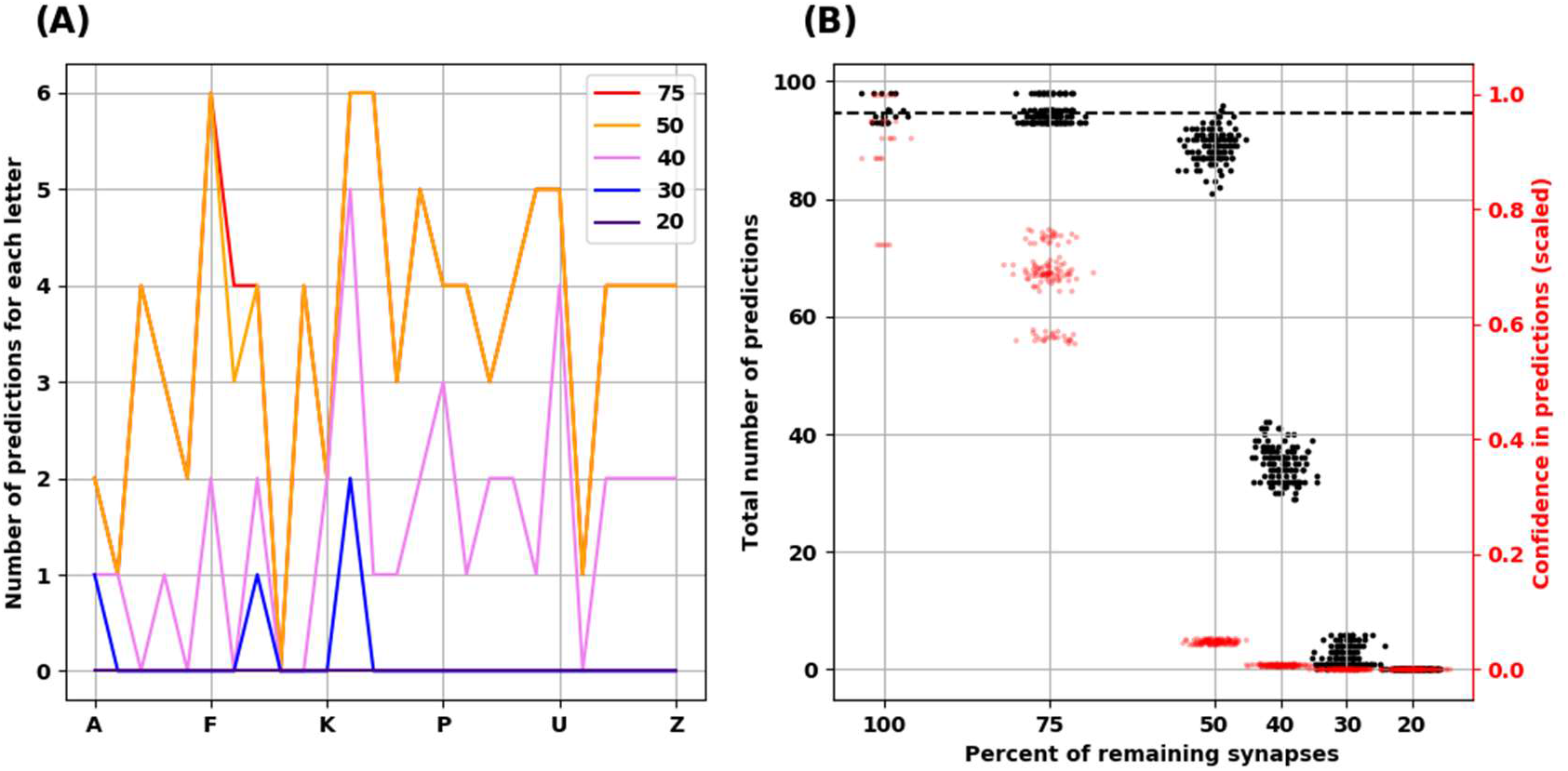
Reduced confidence in predictions before number of predictions is decreased with destroying synapses. **(A)** Illustrative example from one network showing the progressive reduction in number of predictions for each letter with loss of synapses. Each colored line represents the number of remaining synapses. Of note, the number of predictions from network with remaining 75% is identical to network with 100% of synapses. **(B)** At 75% of synapses, the networks encode similar number of predictions as 100% of synapses, but with less confidence. At 50% of synapses, there is a slight reduction of number of predictions, but the confidence is less than 0.1. There is marked reduction of number of predictions at 40 and 30% of synapses. No predictions could be generated with 20% of synapses. There are 25 simulations at 100%, and 125 simulations at the remaining percentages.

The reduction we found in the number of predictions with greater degrees of synaptic loss was not accounted for by less activity in the network. On the contrary, with increasing synaptic destruction, more cells were firing as more minicolumns became fully active. We calculated the average confidence the network had with the progressive destruction of synapses by calculating a scaled ratio of winner cells to active cells (Fig. 4B, red data points and right y-axis). As more synapses were lost, fewer neurons received the predictions from the active neurons on the previous time step, resulting in more bursts of the minicolumns, denoting prediction error. Thus, the less confident the network is, the more bursting and the more active it is. Interestingly, we found that network confidence dropped much faster than its total number of predictions. With 100% of synapses present, the mean confidence was 0.9 (std. dev. = 0.1), while with 75% of synapses remaining, the mean confidence dropped to 0.68 (std. dev. = 0.06) despite the same number of predictions made. With 50% reduction of synapses, despite a mild reduction in the number of total predictions, the confidence dropped to 0.05 (std. dev. = 0.003).

## Discussion

The main goal of this study was to mechanistically connect the reduction of synapses seen in postmortem biopsies from MDD patients and in animal models of depression with MDD symptomatology based on the Bayesian brain, predictive coding, and interoception frameworks. Using the temporal memory (TM) model, we ran computer simulations investigating how the prediction making capacity of a neocortical layer would be affected by reducing the number of available synapses since synaptic loss characterizes brains with MDD and restored number of synapses is associated with antidepressant effects.

Emotions are hypothesized to exist on two dimensions (32). The first is valence, which reflects a prediction about net energy balance. A prediction of negative net energy balance, either because of predicted reduced gain or predicted high energy cost, will be perceived as sadness. The second dimension is arousal, which reflects increased prediction error, and is a signal for learning and updating the internal model. MDD was conceptualized as a disorder where interoceptive neocortical regions are stuck in fewer affective states with high prediction errors (32, 40, 60). We started from the first principles of individual neuronal functioning formalized into a functional microcircuit of a single neocortical layer in the HTM framework (52), and we found that reducing the number of synapses would result in *both* a limited number of predictions and increased prediction error.

Reducing the synapses by 50% slightly reduced the number of predictions made, but the confidence strikingly dropped. Reducing the synapses by more than 50% decreased the number of predictions made by the trained networks further in a non-linear fashion. The decrease in prediction numbers was not due to reduced activity following reduced synaptic numbers. On the contrary, neuronal firing increased secondary to increased minicolumn bursts, reflecting higher prediction error (61) (or lower confidence in predictions).

### Clinical relevance & proposed mechanisms

Rumination, where the brain keeps coming back to the same thought content (62), is construed here as a manifestation of limited predictions and reduced confidence in these predictions. Being stuck in a negative mood could be considered “affective rumination,” or more formally, affective inertia (63). Koval et al. (62) found that the symptom of being cognitively stuck (i.e., rumination) and emotionally stuck (emotional inertia) are positively correlated with one another yet independent in determining the overall severity of the clinical depression. According to the Embodied Predictive Interoception Coding (EPIC) model, the brain’s interoceptive agranular cortices (39) project diffusely to other cortical areas to relay affective states (64, 65). Applying our results to the agranular cortices, e.g., subgenual cingulate cortex (sgACC), suggests a mechanism by which a condition like depression with reduced synapses has limited interoceptive predictive states and high prediction errors, reflecting limited mood experiences, i.e., constricted/restricted affect but with high arousal and perhaps heightened anxiety levels (66, 67), shaping further expectations of patients (68).

Our study also suggests why the limited predicted states in MDD are experienced as negative affective states. According to the concept of interoceptive allostasis, states with predictions of negative energy balance would be perceived as affectively unfavorable (40). When there is reduced confidence in predictions made by interoceptive cortices, the uncertainty would be transmitted to other neocortical and subcortical regions (69, 70). Such uncertainty might signal the need to be “ready for anything,” to prepare for a larger number of possibilities. However, being ready for anything would also predict high energy costs and negative energy balance, which would be perceived by the depressed patient as negative affect.

The non-linear relationship of our findings could explain the rapid antidepressant properties of ketamine and psilocybin. Above a threshold, restoring a fraction of synapses would be enough to improve the confidence in the predictions made, reducing the prediction error signal, thus limiting the number of possibilities for which energy costs have to be calculated, and ultimately reducing the negative energy balance.

Our findings also suggest a possible mechanism for how atrophy in cortical regions in MDD (71–74) could be associated with increased neuronal activity. Multiple studies showed a spontaneous state of hypermetabolism of sgACC in depression, particularly in patients struggling with treatment-refractory depression (75) and as suggested by human MEG studies (76). This is similar to observations arising in mouse models of depression (77). Our results shows that synaptic and dendritic atrophy would lead to less neurons in predictive states, resulting in more columnar bursting, and thus increased neuronal firing.

This work illustrates how to formalize Bayesian brain theories into a neocortical neurobiological framework. For example, suicidal ideations can result from a limited ability to consider alternative future scenarios to the extent that the patient cannot escape internal pain, a construct called internal entrapment. De Beurs et al. (78) investigated the interactions of two prominent theories underlying suicidality, namely interpersonal psychological theory (IPT) and the integrated motivational-volitional (IMV) model. They found that two factors, one from each theory, were directly related to suicidal ideations; the factors were *perceived burdensomeness* of IPT and *internal entrapment* from the IMV framework. In light of the current findings, perceived burdensomeness, viewed as difficulty updating a perceived state, would be related to the restricted ability to make new predictions secondary to the limited number of synapses. Internal entrapment describes the inability to escape internal pain, which could be conceptualized as a limited view of new possibilities because of “tunnel vision” (79). Such a limited view would be expected when motivation visceromotor cortices are unable to make more predictions because of a reduced number of spines and/or decreased spine maturation, a finding supported by studies of BDNF in postmortem brains of people who died by suicide (80, 81). In fact, the notion that functional synaptic connection may be the final common pathophysiologic mediator of depression (82) is consistent with our synapse model presented here, and is also suggested by the following:

1. Stress-based animal models induce BDNF methylation resulting in decreased expression, while antidepressants renew BDNF production through acetylation (83).
2. Likewise, in humans, decreased BDNF serum levels are seen in depressed patients, and these levels correct to the level of healthy controls with selective-serotonin reuptake inhibitors (SSRIs) (84).
3. Synaptic density is decreased in patients with depression, detected by positron emission tomography (PET) (85), while BDNF induces dendritic arborization (86) and neurogenesis, which underlies reversal of depressive behavior in animal models after 1-month of SSRIs (87) or tricyclic antidepressants (88).
4. Electroconvulsive therapy also enhances BDNF production, spine maturation, and neurogenesis (15, 16).

Therefore, synapses and BDNF are reduced in depression, but restoration of BDNF rescues synaptogenesis, reversing depression.

### Relation to other models

Other computer modeling work of MDD has shown how cortical regions can be caught in their dynamic states in MDD. A computer model was developed of two interacting regions, ventral anterior cingulate cortex (vACC) engaged in emotional processing, and DLPFC involved in cognitive processing. Ramirez-Mahaluf et al. used the model to investigate how the two regions interacted together in depression (89). Each region contained 800 pyramidal neurons and 200 interneurons, reciprocally connected. The two regions inhibited each other by activating the interneuronal population in the other region. They modeled a healthy condition where each region can either be active in a state of low-rate asynchronous firing (0.5 - 1 spikes/second) or high-rate asynchronous firing (25-30 spikes/second). The corresponding region became active in the healthy network when presented with either a cognitive (working memory) or an emotional (sadness provocation) task. When they modeled depression as slower glutamate reuptake in vACC, it resulted in sustained activity of vACC that DLPFC could not inhibit. Thus, the network could not switch to active DLPFC when presented with the cognitive task. They explored the progression of MDD symptom severity from mild to moderate to severe, and found that switching to active DLPFC functioning in the face of a cognitive task was more difficult as the disease progressed. They examined the effect of SSRIs on the dynamics of switching and found that SSRIs inhibited the excitatory neuronal population in the vACC and reversed the dynamics back to the healthy bistable state, though the reversal became more difficult in more severe depression. When they modeled the effects of deep brain stimulation (DBS) on their model, they found that DBS also reversed the dynamics. Similar to our study, modeling depression resulted in increased activity of the interoceptive region model (vACC), where the increased firing was secondary to increased prediction errors. Somatostatin-positive interneurons (SST) are a population of interneurons that mainly target the distal apical dendrites of pyramidal neurons in the neocortex. Deficiency of SST interneurons might play a role in MDD (90). To examine the effects of reducing somatostatin (SST) mRNA in postmortem tissue from depressed patients (91), Yao et al. (92) developed a computer model of neocortical layer 2/3 based on both human and rodent data. The model consisted of pyramidal neurons targeting three interneuronal populations: SST, parvalbumin-positive (PV), and vasoactive intestinal peptide-positive (VIP) interneurons. SST interneurons targeted the apical dendrites of pyramidal neurons, as well as the PV and VIP interneurons. To model depression, they decreased the SST inhibition by 40% and examined how that affected neuronal firing and stimulus processing. SST activity reduction increased the background firing rates of the different neuronal populations, including the pyramidal neurons. However, when they presented a stimulus to their model, the stimulus triggered similar pyramidal neuronal firing rates. Similar to our findings, the signal-to-noise ratio in that model (calculated as the ratio between baseline pyramidal firing rate and the post-stimulus firing rate) was reduced because the baseline firing rate was higher in the depressed network model than in the healthy network model. They also found that reduction in the signal-to-noise ratio doubled the rates of both failed detection and false detection of the stimulus when it was presented in the depressed models.

In the light of evolutionary psychology and the Bayesian brain framework, one theory conceptualizes depression as a way to minimize surprises in social interactions (93). This theory is based on observations that patients diagnosed with depression have learned to expect negative interactions with others based on their past experiences. This would reduce surprises (i.e., prediction error), perhaps as a mechanism to reduce emotional vulnerability when such negative interactions arise. At the same time, this theory assumes that it is challenging for a depressed patient to update their (negatively-valenced) beliefs. Given their predictions about negative interactions, it is possible that depressed patients tend to avoid social interactions, and thus miss interactions that could update their expectations about the outcome of the social interaction. Another possibility could be that neuromodulatory neurobiological factors in a depressed brain, such as reward learning mediated by dopamine neurotransmission, make it difficult for patients to update their predictions.

The findings from our study hint at a third potential mechanism underlying the difficulty a depressed patient may have in updating their negative cognitions. Our results show that a reduced number of synapses to encode more predictions will limit the ability to update one’s beliefs with novel interactions or experiences. Typically, authors of prior studies with Bayesian brain formulations of psychiatric disorders (e.g., (93, 94)) have assumed that a distinct neuronal population be used for calculating predictions than the population used for calculating prediction errors. Barrett et al. (39) suggested that agranular layers of interoceptive cortices can send predictions but lack the number of neurons needed to compute prediction errors of interoceptive inputs. In the TM model of the HTM framework that we used here, neurons within a single neocortical layer were capable of both making predictions and calculating prediction errors. This suggests that agranular cortical areas can both send predictions and compute prediction errors.

### Future Directions

This is a proof-of-concept study of a single neocortical layer exhibiting synaptic atrophy as seen in MDD and animal models of depression. David Marr suggested three levels of analysis when looking at the function of neuronal microcircuits (95, 96). The first is a computational level: what is the problem that the microcircuit solves? The second level is the algorithmic level, describing the steps needed to solve the problem. The third level is the implementation level, how the microcircuit elements, such as neuronal populations and their connections, perform the steps of the algorithm. We aimed to connect Marr’s three levels of analysis in the context of MDD using the HTM framework. At the computational level, the problem is making predictions about energy expenditures in a changing environment. At the algorithmic level, the network represents multiple possible predictions and selects the ones which agree with the current input (52). At the implementation level, the neurons used are restrained by the neurobiology of neocortical neurons, e.g. sub-threshold soma depolarization by NMDA spikes when clustered synapses are activated within a short time window (55, 97).

To validate the main prediction from this work, we can look at sequences of action potentials, or replays. According to the HTM framework, predictions are encoded as temporal firing sequences. Replay of neocortical temporal sequences has been shown widely across multiple neocortical regions, usually in the context of memory retrieval (98, 99), as well as in neocortical slices (100). Replay could be examined in animal models of depression, and from invasive recordings in patients undergoing DBS for depression, as in Scangos et al. (101). We predict that the variability of the replays will be reduced in depression.

There are several ways to take this work further. The dendrites represented here are basal. Within the structural model, agranular cortical areas receive input from dysgranular cortical areas (with less developed layer IV) onto deeper layers (102), most likely terminating on basal dendrites of layer 5 and 6 neurons. Both ketamine (103) and psilocybin (31) increased the number of spines along the basal dendrites, as well as along the apical dendrites (e.g. (26, 30)). Adding apical dendrites to the model will allow us to examine how apical synaptic atrophy affects the predictive function of the neocortex, especially with the postulated role of SST interneurons targeting apical dendrites in depression (91).

Another future direction involves expanding the model to include multiple HTM layers. We can then investigate the effect of loss of specific synapses on predictions made by a whole cortical column (104), and the effects of synapses within layers versus between layers. Yet another future direction would be to implement neuromodulators like monoamines that play a role in depression. Our model does not take the effects of endogenous neuromodulators such as serotonin or dopamine into account. The level of modeling we did here also does not consider the immunological and hormonal mechanisms that mediate the effects of chronic stress on synaptic atrophy. Instead, we looked at one of the possible outcomes of chronic stress, namely synaptic atrophy, and how this could be related to MDD.

To our knowledge, this is the first study applying algorithms from the biologically constrained HTM framework to neocortical abnormalities found in a psychiatric disorder. This work highlights the potential of the HTM framework to formalize the predictive capacity of neocortical layers and the possible insights this line of investigation can bring to understanding psychiatric disorders. Instead of limiting brain microcircuit modeling to molecular and microcircuit mechanisms underlying specific neural dynamics, this approach extends microcircuit modeling into the functional realm, potentially connecting microcircuit models to behavioral tasks and symptoms (105–107), using neurons that have more biological features than ones traditionally used in artificial neural networks.

## Acknowledgments and conflicts of interest

MAS is a consultant for *In Silico* Biosciences, Inc. LLC has consulted for Otsuka, Sunovion, Neuronetics, Nexstim, Janssen, Affect Neuro, Sage Therapeutics, and Neurolief. She has research support through clinical trial contracts at Butler Hospital with Janssen, Neuronetics, Affect Neuro, and Neurolief. The other authors has nothing to declare. MAS is supported by the National Institute of Mental Health (NIMH) (R21MH125199). JCB is supported by the National Institute of General Medical Sciences (NIGMS) (P20GM130452). Graphical abstract was created by BioRender.com.

